# Hydroxylation site-specific and production-dependent roles of endogenous oxysterols in cellular cholesterol homeostasis

**DOI:** 10.1101/2022.05.14.491815

**Authors:** Hodaka Saito, Ryuichiro Sato, Makoto Shimizu, Yoshio Yamauchi

**Author notes:** Corresponding author: Yoshio Yamauchi, Ph.D., Department of Applied Biological Chemistry, Graduate School of Agricultural and Life Sciences, The University of Tokyo, 1-1-1 Yayoi, Bunkyo-ku, Tokyo, Japan. **Author contributions:** Y.Y. conceived research. H.S., R.S., and Y.Y. designed experiments. M.S. contributed reagents. H.S., R.S., and Y.Y. analyzed data. H.S. and Y.Y. wrote manuscript.

## Abstract

The cholesterol metabolites oxysterols play central roles in cholesterol feedback control. They modulate the activity of two master transcription factors that control cholesterol homeostatic responses, sterol regulatory element-binding protein-2 (SREBP-2) and liver X receptor (LXR). Although the role of exogenous oxysterols has been well-established, whether endogenously synthesized oxysterols similarly control both SREBP-2 and LXR to those added exogenously remains poorly explored. Here, we carefully validate the role of oxysterols enzymatically synthesized within cells in cholesterol homeostatic responses. We first show that SREBP-2 responds more sensitively to exogenous oxysterols than LXR. We then show that endogenous 25-hydroxycholesterol (25-HC), 27-HC, and 24S-HC synthesized by CH25H, CYP27A1, and CYP46A1, respectively, suppress SREBP-2 activity at different degrees by stabilizing Insig proteins whereas 7a-HC has little impact on SREBP-2. The results explain the hydroxylation site-specific role of endogenous oxysterols. On the other hand, the expression of CH25H, CYP46A1, CYP27A1, or CYP7A1 fails to induce LXR target gene expression. We also show the 25-HC production-dependent suppression of SREBP-2 using a tetracycline-inducible CH25H expression system. Moreover, we quantitatively determine the specificity of the four cholesterol hydroxylases in living cells. Based on these results, we propose that endogenous side-chain oxysterols primarily regulate the activity of SREBP-2, not LXR.

## Introduction

Cholesterol homeostasis is strictly controlled by multiple feedback mechanisms (1–3), and its dysregulation is associated with diverse pathophysiological settings from congenital to acquired human diseases (4–6). Not only cholesterol but also other sterol molecules, including oxysterols, participate in the feedback controls. The cholesterol metabolites oxysterols serve as crucial mediators that regulate cellular cholesterol homeostasis (7). In particular, oxysterols whose side-chain is hydroxylated (hereafter referred to as side-chain oxysterols) exert much more potent effects than sterol-ring oxysterols. The oxysterol-mediated control of cholesterol homeostasis involves two distinct transcription factors; sterol regulatory element-binding protein-2 (SREBP-2) and liver X receptor (LXR). SREBP-2 regulates the expression of most enzymes involved in cholesterol biosynthesis and low-density lipoprotein receptor (LDLR) that mediates cholesterol uptake from the extracellular milieu (8, 9). On the other hand, the nuclear receptor LXR regulates the removal of excess cellular cholesterol by enhancing the expression of several ABC transporters that export cholesterol from cells, including ABCA1 and ABCG1 (6, 10).

SREBP-2 is a membrane-bound transcription factor. As an inactive form, SREBP-2 resides in the endoplasmic reticulum (ER), where it forms a complex with SREBP cleavage activating protein (SCAP) and Insig proteins (9). SCAP serves as an SREBP-2 chaperone for its transport to the Golgi, while Insig acts as an ER retention factor for SREBP-2 (1). For the activation, the SREBP-2/SCAP complex dissociates from Insig and is transported to the Golgi for the proteolytic cleavage and release of the transcriptionally active domain. Its transport to the Golgi is tightly regulated by cholesterol and oxysterols. Cholesterol binds to SCAP, which leads to its conformational change for blocking the incorporation of the SREBP-2/SCAP complex into the COP-II vesicles (11, 12), while oxysterols interact with Insig proteins and prevent their proteasomal degradation (13). These events cooperatively prevent the exit of SREBP-2 from the ER and thus negatively regulate cholesterol biosynthesis and uptake (1). Another oxysterol effector, LXR exhibits high-affinity binding to various oxysterols (14). When added to the culture medium, oxysterols bind to LXR as its ligand and activate its transcriptional activity. LXR forms a heterodimer with another nuclear receptor, retinoid X receptor (RXR), and transactivates the target genes to promote cellular cholesterol efflux.

Cholesterol undergoes oxygenation through enzymatic and non-enzymatic processes, and a variety of sterol hydroxylases catalyze the synthesis of oxysterols (15, 16). Although a number of oxysterols are identified in the circulation and tissues, the major oxysterols found in the body are 7α-hydroxycholesterol (7a-HC), 27-HC, 24S-HC, and 25-HC. These four oxysterols are largely synthesized by CYP7A1, CYP27A1, CYP46A1, and CH25H, respectively, in the ER or mitochondria (16, 17). 27-HC, 24S-HC, and 25-HC are all capable of regulating both SREBP-2 and LXR when added to cells exogenously. On the other hand, 7a-HC, a major bile acid precursor, elicits only subtle effects on these two effectors.

As such, it is well-established that oxysterols have dual roles in cholesterol homeostasis: regulation of cholesterol acquisition (biosynthesis and uptake) and cellular cholesterol removal. However, there has been controversy regarding the role of oxysterols in cholesterol homeostasis (7, 18). One of the reasons is that the current knowledge on oxysterol-mediated regulation is primarily based on experimental utilization of supraphysiological levels of oxysterol added exogenously to cells (hereafter referred to as “exogenous” oxysterols), despite the evidence that oxysterols are present in the circulation and tissues at low levels (15, 19). Little is known on whether endogenously synthesized oxysterols (hereafter referred to as “endogenous” oxysterols) elicit the same effects as exogenous oxysterols on cellular cholesterol homeostasis. To address these issues, we seek to explore whether endogenous oxysterols also play dual roles in cellular cholesterol homeostasis like exogenous oxysterols by expressing the four major cholesterol hydroxylases CH25H, CYP27A1, CYP46A1, and CYP7A1 in wild-type and mutant Chinese hamster ovary (CHO) cells that have extensively been utilized for studying cholesterol homeostatic responses. We also aim to determine the specificity of these enzymes quantitatively in living cells.

## Results

### SREBP-2 is more sensitive to exogenous side-chain oxysterols than LXR

We selected four major oxysterols enzymatically synthesized in the body, 7a-HC, 24S-HC, 25-HC, and 27-HC (**Figure 1A**). Minimum amounts of each oxysterol required for modulating SREBP-2 and LXR activities are not fully characterized in a single cell system. Therefore, we first carefully validated dose-dependent effects of these four oxysterols on cholesterol homeostatic responses in CHO-K1 cells. To determine the dose-dependence of oxysterols added exogenously, the cells were treated with 7a-HC, 24S-HC, 25-HC, or 27-HC at different concentrations. The results show that all the side-chain oxysterols suppress the expression of SREBP-2 target genes *(Hmgcs1, Hmgcr, Sqs,* and *Lss)* in a dose-dependent manner, but their inhibitory potencies are dependent on hydroxylation sites (**Figure 1B, Figure S1A**). In particular, 25-HC has the most potent inhibitory effect on the expression of the SREBP-2 target genes; it inhibits *Hmgcs1* expression at as low as 20 nM. 27-HC and 24S-HC also suppressed the expression of SREBP-2 targets genes when added at 100 nM, but their inhibitory effects were lower than 25-HC. In contrast to the repression of SREBP-2 target genes at the low concentrations, 25-HC, 27-HC, and 24S-HC failed to increase the expression of the LXR target genes *(Abca1* and *Abcg1)* at 100 nM or lower **(Figure 1B and Figure S1B**). Five hundred nM or higher concentrations of oxysterols were required for up-regulating the expression of LXR target genes. On the other hand, 7a-HC showed no effects on mRNA levels of SREBP-2 and LXR target genes even treated at 2.5 μM. These results indicate that side-chain oxysterols preferentially down-regulate SREBP-2, whereas they activate LXR at higher concentrations without adding the RXR ligand *9-cis* retinoic acid.

**Figure 1.**
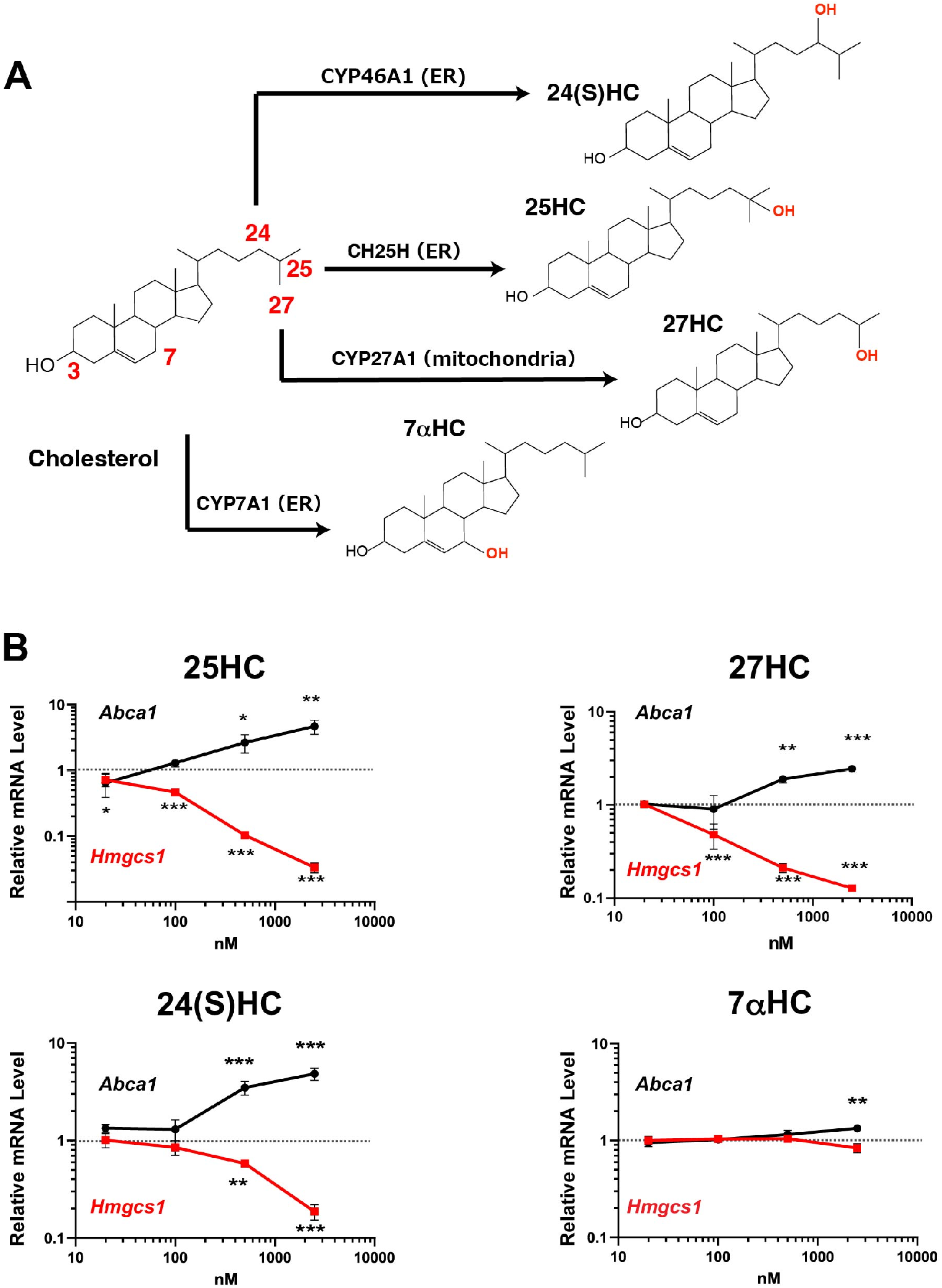
SREBP-2 is more sensitive to exogenous side-chain oxysterols than LXR. (A) Biosynthesis of oxysterols by cholesterol hydroxylases, CH25H, CYP27A1, CYP46A1, and CYP7A1. (B) CHO-K1 cells were treated without or with 25-HC, 27-HC, 24S-HC, or 7a-HC at different concentration (0, 20, 100, 500, and 2500 nM) for 24 hr. mRNA levels of SREBP-2 and LXR target genes *(Hmgcs1* and *Abca1,* respectively) were analyzed by qPCR. Data represent means ± SD (n = 3). Statistical analyses were performed by one-way ANOVA with Dunnett post hoc test by comparing to the vehicle treatment group (*p<0.05, **p<0.01, ***p<0.001).

### Cholesterol hydroxylase expression modulates cholesterol homeostatic responses

Circulating oxysterol levels are as low as (lower than) approximately 30 – 150 ng/ml (80 - 400 nM), which is 0.001 – 0.01% of circulating cholesterol (15). Although 25-HC, 27-HC, 24S-HC, and 7a-HC are all enzymatically synthesized from cholesterol within cells, the effects of cholesterol hydroxylase expression on cholesterol homeostatic responses remain poorly characterized at cellular levels. Therefore, we sought to explore how the expression of the major cholesterol hydroxylases, CH25H, CYP27A1, CYP46A1, or CYP7A1, alters cholesterol homeostatic responses in CHO-K1 cells. We first examined the cellular localization of CH25H, CYP27A1, CYP46A1, and CYP7A1 forcedly expressed. The results show that CH25H, CYP46A1, and CYP7A1 reside in the ER whereas CYP27A1 localizes to the mitochondria, confirming their correct localization (**Figure 2A**). Furthermore, CH25H, CYP46A1, and CYP7A1 partly colocalize with the mitochondria marker TOM20, suggesting the localization of these enzymes at the ER-mitochondria contact site known as mitochondria-associated membranes (MAMs) (**Figure S2A**). Next, we examined the effect of hydroxylase expression on SREBP-2 activity. The results showed that the expression of CH25H, CYP27A1, and CYP46A1 markedly decreased mRNA levels of SREBP-2 target genes *(Hmgcs1, Hmgcr, Sqs,* and *Lss),* whereas CYP7A1 expression showed only subtle effects on the expression of these genes (**Figure 2B**). Consistent with these results, the expression of CH25H, CYP27A1, and CYP46A1 reduced the *Hmgcs1* promoter activity as much as the addition of the corresponding oxysterols (**Figure S2B**). These results also suggest that the expression of these hydroxylases produces significant amounts of corresponding oxysterols. The effect of CH25H expression on SREBP-2 was dependent on cholesterol 25-hydroxylase activity as the mutant CH25H^H242Q/H243Q^ substantially lacking the hydroxylase activity (20) was much less efficient in repressing *Hmgcs1* and *Hmgcr* expression (**Figure S2C**). Furthermore, it was shown that in HEK293T cells where the mitochondrial cholesterol transporter StAR (also known as STARD1) is lacking, StAR was required for suppressing mRNA levels of SREBP-2 target genes by CYP27A1 expression (**Figure S2D**), confirming a previous report (21). We also monitored the expression of LXR target genes in cells overexpressing these four hydroxylases. In contrast to exogenous oxysterols at 2.5 μM enough high for induction, the expression of hydroxylases failed to increase mRNA levels of *Abca1* and *Abcg1* (**Figure 2C**).

**Figure 2.**
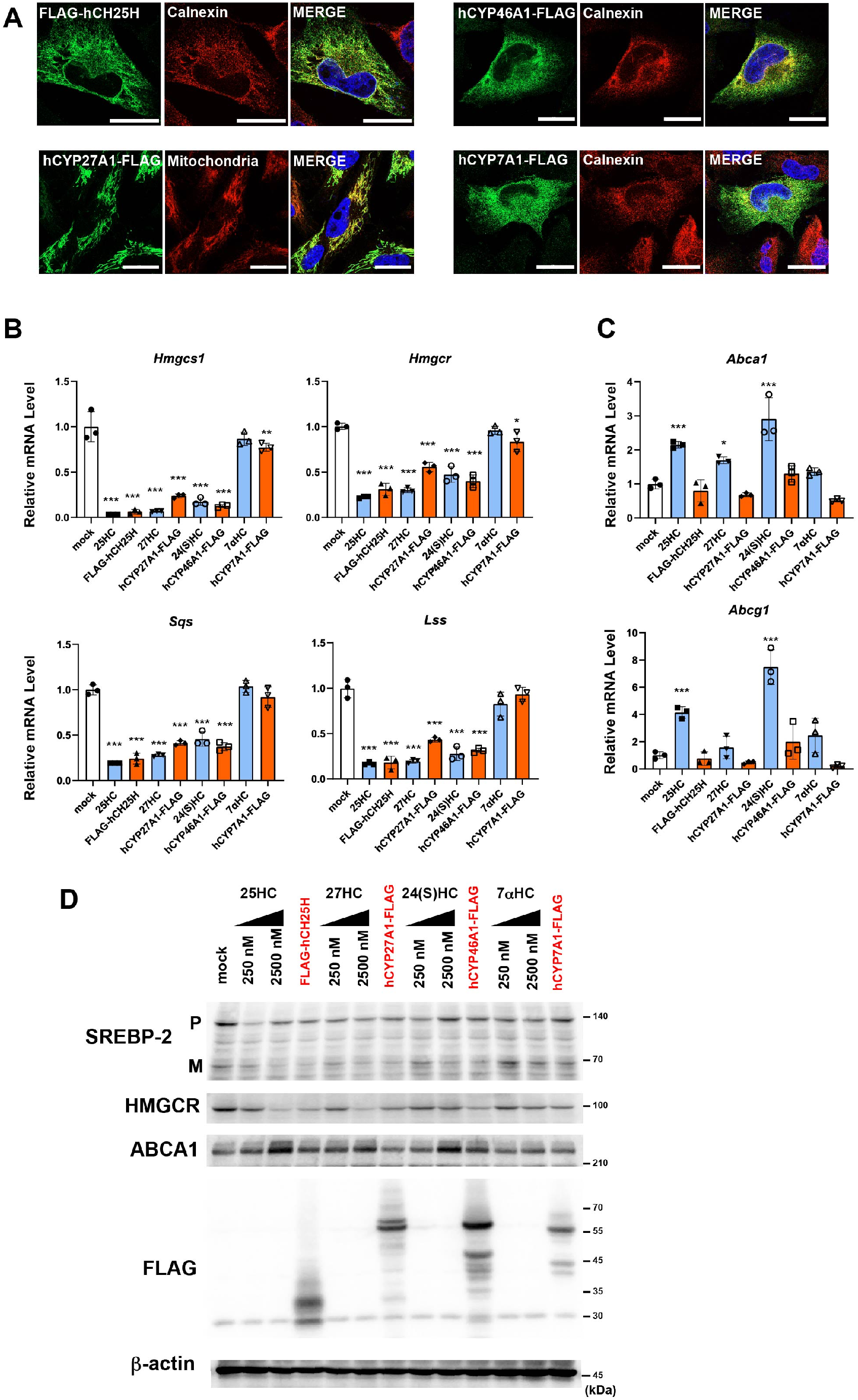
Effect of cholesterol hydroxylase expression of cholesterol homeostatic responses. (A) Cellular localization of cholesterol hydroxylases. A2058 cells were transfected with the plasmids (pFLAG-CH25H, pCYP27A1-FLAG, pCYP46A1-FLAG, or pCYP7A1-FLAG) and fixed 2 days after transfection. Cholesterol hydroxylases were labeled with anti-FLAG antibody followed by Alexa 488-conjugated anti-mouse IgG (green). The ER and mitochondria were labeled with anti-calnexin and anti-Tom20 antibodies, respectively, followed by Alexa 568-conjugated anti-rabbit IgG (red). Images were taken under a confocal microscopy as described in Materials and Methods. Scale bar, 20 μm except FLAG-CH25H (10 μm). (B and C) Effect of hydroxylase expression on cholesterol homeostatic responses. CHO-K1 cells were transfected with plasmids as above and incubated for 48 hr in medium containing 0.1% FBS. Cells transfected with pCMV10 (mock) were also treated without or with 2.5 μM oxysterols (25-HC, 27-HC, 24S-HC, or 7a-HC) for 16 hr in medium containing 0.1% FBS before harvesting cells. mRNA levels of SREBP-2 target genes (B) and of LXR target genes (C) were analyzed by qPCR. Data represent means ± SD (n = 3). Statistical analyses were performed by one-way ANOVA with Dunnett post hoc test by comparing to mock (*p<0.05, **p<0.01, ***p<0.001). (D) Effect of hydroxylase expression on SREBP-2 processing. CHO-K1 cells were transfected with the plasmids and treated as above. Before harvesting, cells were incubated with MG132 (20 μM) for 2 hr. Expression of SREBP-2 (P, precursor; M, mature form), HMGCR, ABCA1, and FLAG-tagged hydroxylases was analyzed by immunoblot. β-actin serves as a loading control.

To further explore the effects of the hydroxylase expression on cholesterol homeostatic responses, we assessed SREBP-2 processing and HMGCR and ABCA1 protein expression. As expected, the expression of CH25H, CYP27A1, and CYP46A1, but not CYP7A1, markedly inhibited SREBP-2 processing as strong as exogenous side-chain oxysterols (2.5 μM) did (**Figure 2D**). Consistently, HMGCR protein level was also reduced by CH25H, CYP27A1, and CYP46A1. In contrast to SREBP-2 target genes, expressing these hydroxylases did not increase ABCA1 protein levels while its expression was induced by exogenous 25-HC, 27-HC, and 24S-HC. These results demonstrate that *de novo* synthesized side-chain oxysterols primarily regulate SREBP-2 activity but have little impact on LXR activity.

### 25HC production-dependent inactivation of SREBP-2

Our results described above show that CH25H expression and its product 25-HC have a more potent inhibitory effect on SREBP-2 activity than other hydroxylases and oxysterols. To more carefully examine the impact of CH25H expression, we took advantage of the doxycycline (Dox)-inducible CH25H expression system that enables us to control the levels and timing of its expression. We established CHO-CH25H^tet-on^ cells and examined the effects of CH25H expression on cholesterol homeostatic responses. As expected, Dox induced CH25H expression in a manner dependent on its concentration (**Figure 3A, B**). We first examined the effects of increasing CH25H expression on cholesterol homeostatic responses at mRNA levels. The results showed that the expression of SREBP-2 target genes *(Hmgcs1, Hmgcr, Sqs, Lss)* was suppressed in a manner dependent on *CH25H* expression (**Figure 3A**), but mRNA levels of LXR target genes *(Abca1* and *Abcg1)* were unaltered by the induction of *CH25H* expression. To directly demonstrate the CH25H-dependent repression of SREBP-2, we next examined SREBP-2 processing with an increasing concentration of Dox. We found the inverse correlation between CH25H expression and SREBP-2 processing (**Figure 3B**); the magnitude of reduction in the mature form depended on CH25H expression levels. On the other hand, the increase in ABCA1 protein levels was only subtle even when the cells were treated with Dox at the concentration of 0.8 μg/ml or higher. These data are consistent with the results in Figure 2.

**Figure 3.**
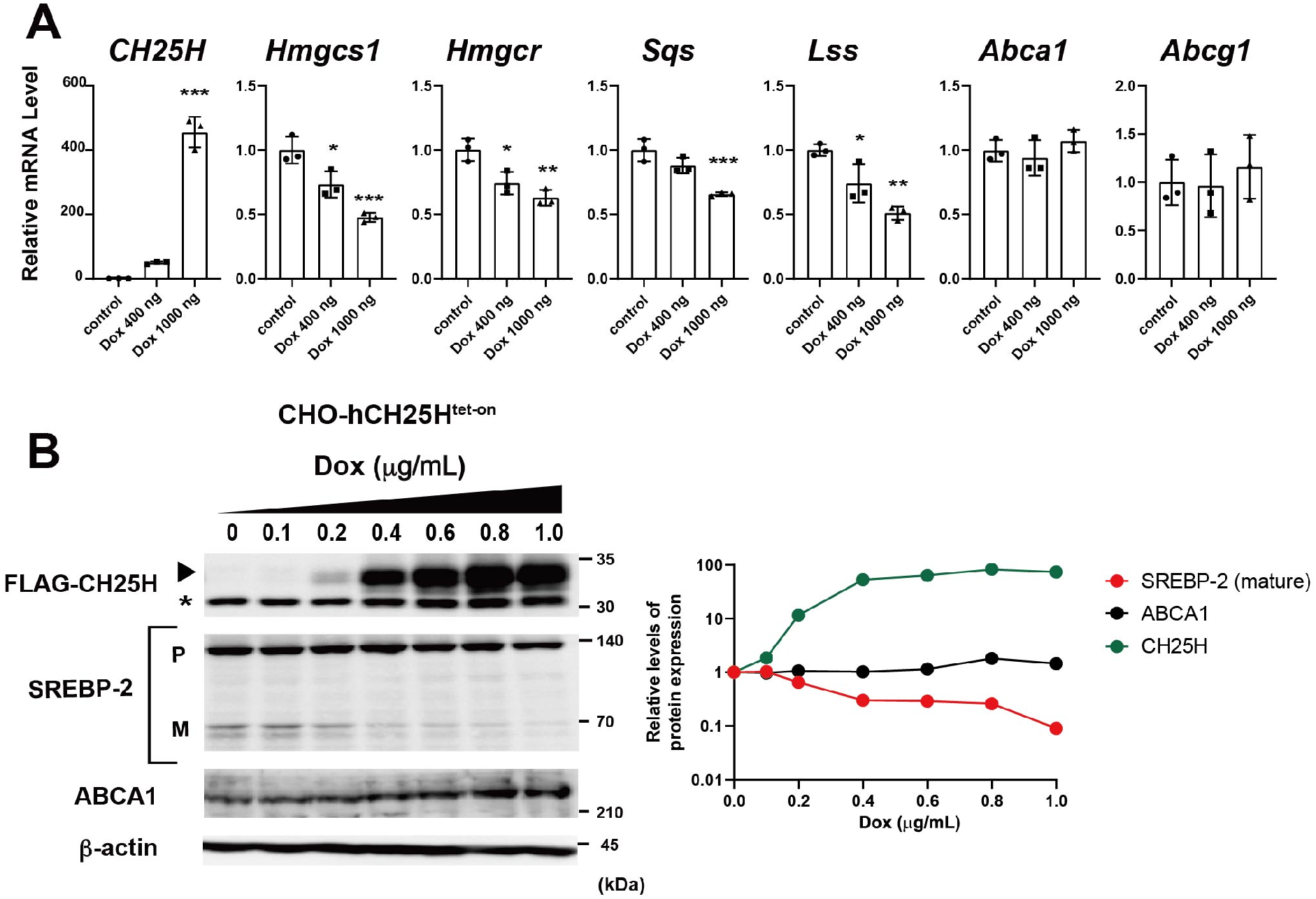
CH25H expression level-dependent inactivation of SREBP-2. (A) Correlation between CH25H expression and SREBP-2 activity. CHO-CH25H^tet-on^ cells were treated without or with 400 and 1,000 ng/mL Dox for 24 hr. mRNA levels of the indicated genes were measured by qPCR. Error bars represent S.D. from three biological replicates. Statistical analyses were performed by one-way ANOVA with Dunnett post hoc test (*p<0.05, **p<0.01, ***p<0.001). (B) Effect of CH25H expression levels on SREBP-2 processing and ABCA1 expression. CHO-CH25H^tet-on^ cells were incubated with different concentration of Dox (0 - 1,000 ng/mL) for 24 hr as indicated. Cell lysate was subjected to immunoblot analysis to detect FLAG-CH25H, SREBP-2 (both precursor (P) and mature (M) forms), and ABCA1. β-actin was detected as a loading control. The asterisk denotes a non-specific band. In the right panel, relative changes in the expression of FLAG-CH25H, SREBP-2 mature form, and ABCA1 are plotted.

### Endogenous side-chain oxysterols regulate Insig-dependent events

The results in Figures 2 and 3 suggest that endogenous side-chain oxysterols selectively regulate SREBP-2 activity. SREBP-2 processing is regulated by SCAP and Insig at the ER by independent mechanisms; Cholesterol binds and regulates SCAP, whereas side-chain oxysterols stabilize Insig by direct interaction. Next, we asked how endogenous oxysterols modulate SREBP-2 activity. To this end, we employed two CHO mutants, 25RA and SRD15, both of which exhibit 25-HC-resistant and constitutive SREBP-2 activation phenotypes through distinct mechanisms (**Figure 4A**); 25RA cells express constitutively active SCAP, namely SCAP^D443N^ (22), while SRD15 cells lack Insig-1 and Insig-2, negative regulators of SREBP (23). We transiently expressed CH25H in these mutant CHO cells and their parental cells and compared the differences in cholesterol homeostatic responses between mutant and parental cells. As previously reported (22, 23), in both 25RA and SRD15 cells, mRNA levels of most SREBP-2 target genes were approximately 2 – 3 folds of their parental cells (**Figure 4B, C**). The expression of CH25H in 25RA cells resulted in a marked reduction in mRNA levels of SREBP-2 target genes in a dose-dependent manner (**Figure 4B, Figure S4A**). However, in 25RA cells forcedly expressed CH25H or treated with 25-HC, these mRNA levels were still higher than parental CHO-K1 cells with the same treatment, thereby providing 25RA cells with 25-HC resistance (22, 23). In sharp contrast to 25RA cells, the reduction in the expression of SREBP-2 target genes upon CH25H expression was much more modest in SRD15 cells (**Figure 4C, Figure S4B**). SRD-15 cells partly retain the response to 25-HC (**Figure S4B**) presumably because the cells express residual levels of Insig-2 (23). These results indicate that endogenous 25-HC suppresses SREBP-2 activity in an Insig-dependent manner.

**Figure 4.**
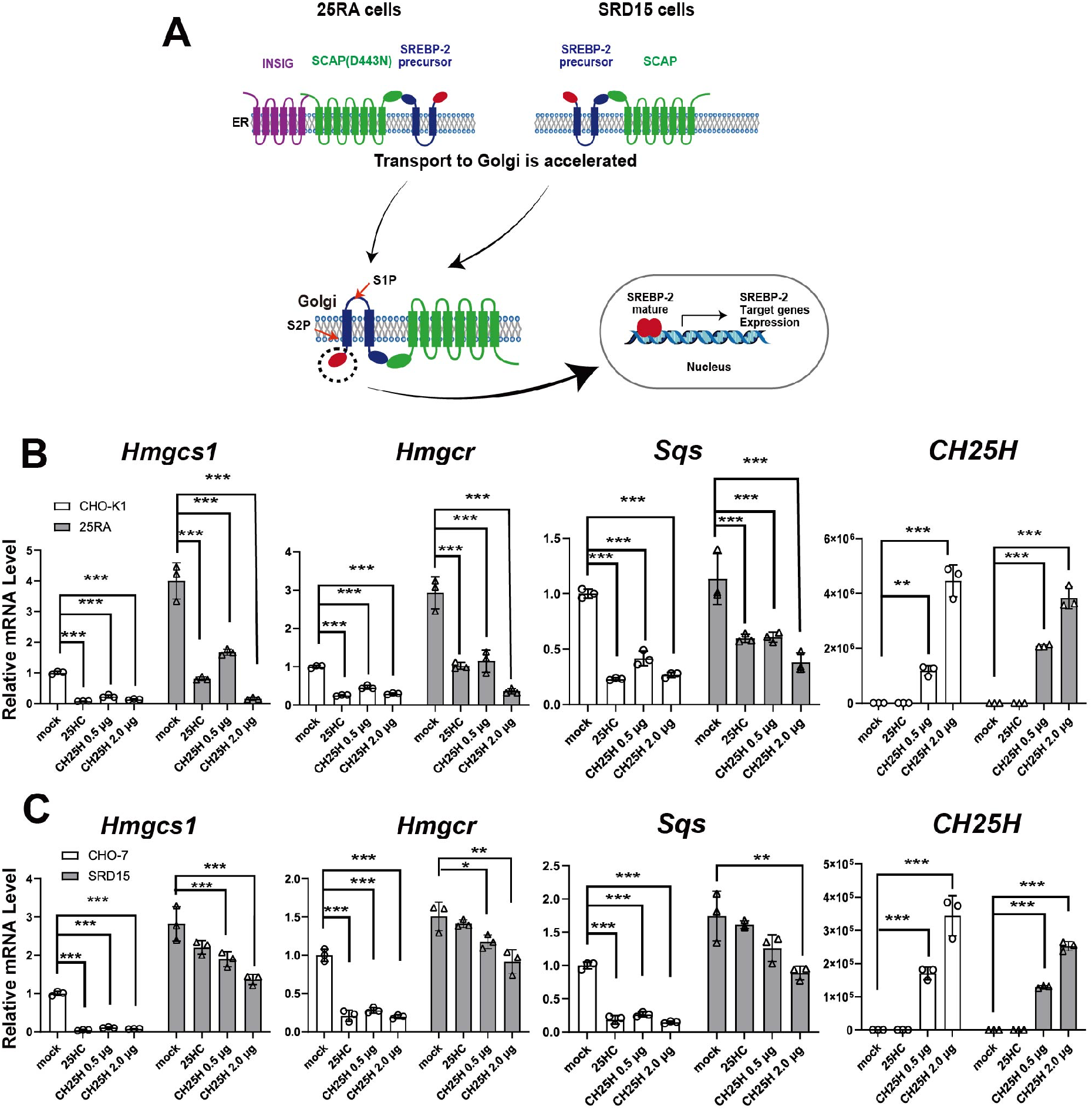
Insigs are required for inactivation of SREBP-2 activity by endogenous oxysterols. (A) Diagram of constitutive activation of SREBP-2 in 25RA and SRD15 cells. In these two mutants, SREBP-2 is constitutively activated by different mechanisms. See text for more detail. (B, C) Expression of SREBP-2 target genes in 25RA (B) and SRD-15 (C) cells. On day 0, 25RA, SRD-15, and their parental cells (CHO-K1 and CHO-7, respectively) were seeded into 6-well plates. On day 1, these cells were transfected with different amounts (0.5 and 2.0 μg/well) of plasmid to express FLAG-CH25H or an empty plasmid as indicated. On day 2, mock-transfected cells were either treated or untreated with 25-HC cells (2.5 μM) for 16 hr. On day 3, cells were harvested for RNA isolation. mRNA levels of SREBP-2 target genes were analyzed by qPCR. Data represent means ± S.D. from three biological replicates. Statistical analyses were performed by one-way ANOVA with Dunnett post hoc test (*p<0.05, **p<0.01, ***p<0.001).

We recently reported that oxysterol-dependent activation of activating transcription factor-4 (ATF4) requires Insig proteins (24). Next, we assessed whether CH25H expression activates the ATF4 axis using the CHO-CH25H^tet-on^ cells. The results show that the induction of CH25H expression was accompanied by the increase in the expression of the ATF4 target genes *Chac1* and *Trb3* in a Dox concentration-dependent manner (**Figure S4C**). Taken together, these results indicate that Insigs are primary mediators for these cellular responses to 25-HC production.

### Endogenous oxysterols stabilize Insig proteins

Oxysterols can bind Insig proteins and protect them from proteasomal degradation, thereby inhibiting SREBP-2 processing (13, 25). We, therefore, sought to directly determine whether endogenous side-chain oxysterols stabilize Insig proteins. The stabilization of Insig was assessed using the translation inhibitor cycloheximide (CHX). As reported, Insig-1 protein levels were rapidly decreased by the CHX treatment in CHO-K1 and HEK293 cells, and the reduction was partly inhibited by adding 25-HC (**Figure 5A, B**). Similar to 25-HC addition, CH25H expression also suppressed Insig-1 degradation, indicating that Insig-1 was stabilized by endogenous 25-HC as efficient as exogenous 25-HC. The stabilization of Insig-1 by CH25H expression was also observed in 25RA cells (**Figure 5A**). Furthermore, the expression of CYP27A1 also inhibited the degradation of Insig-1 in CHO-K1 cells (**Figure S5**).

**Figure 5.**
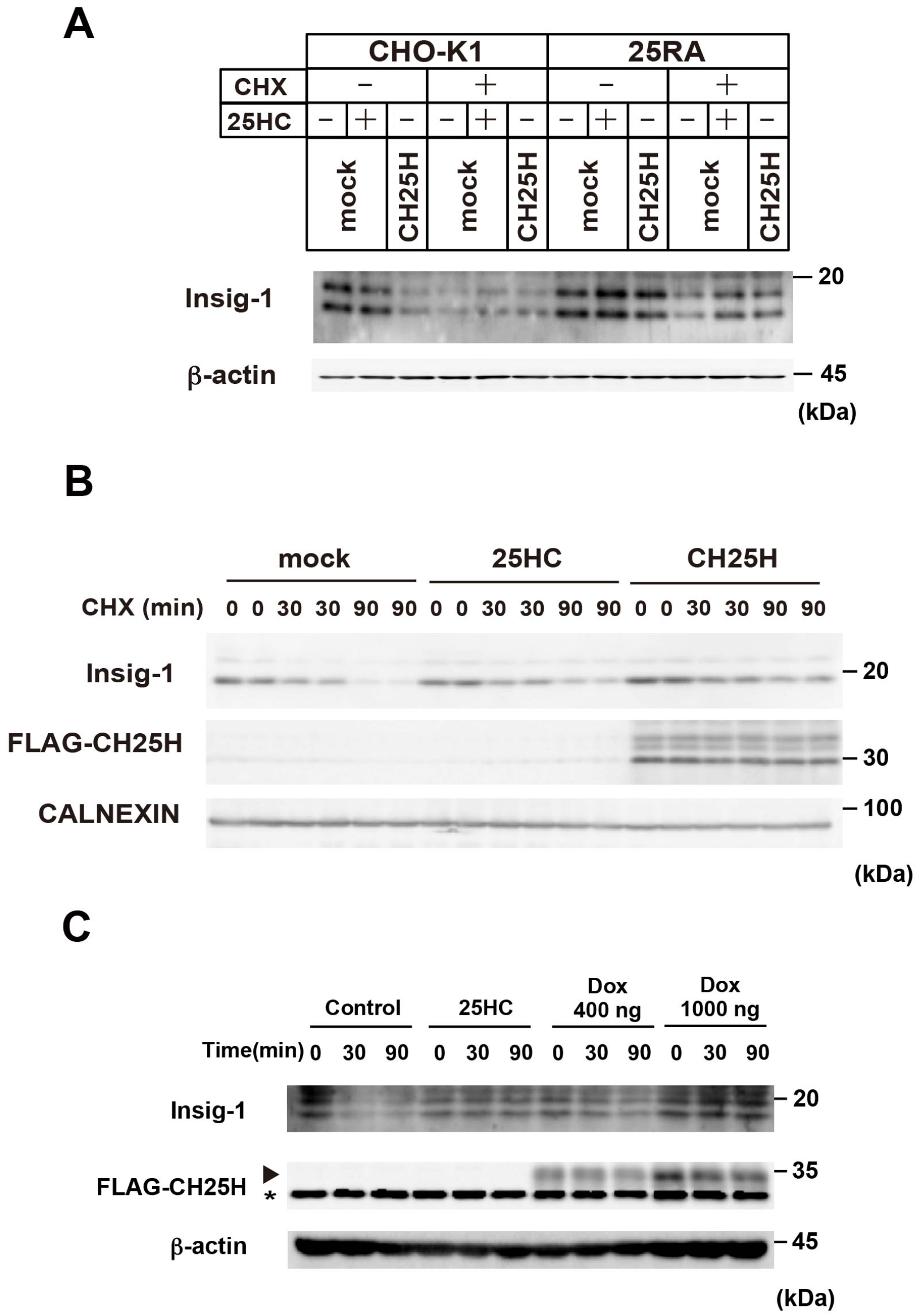
Endogenous oxysterols stabilize Insig-1 proteins. (A) Stabilization of Insig-1 by CH25H expression in CHO cells. On day 1, CHO-K1 and 25RA cells were transfected with plasmid encoding FLAG-CH25H (2 μg/well) as above. On day 2, cells were treated without or with 25-HC (2.5 μM) in the presence or absence of cycloheximide (CHX) (50 μM) for 2 hr as indicated. Immunoblot was performed with antibodies to Insig-1, FLAG, and β-actin. (B) Stabilization of Insig-1 by CH25H expression in HEK293T cells. HEK293T cells were transfected with pFLAG-CH25H (1.0 μg/well) as above. Cells were then treated without or with CHX (50 μM) and 25HC (2.5 μM) for 30 or 90 min as indicated. Stabilization of Insig-1 was examined by immunoblot. Calnexin was used as a loading control. (C) Expression level-dependent effect of CH25H on Insig-1 stabilization. CHO-CH25H^tet-on^ cells were incubated with or without Dox (400 ng/mL or 1000 ng/mL) for 24 hr, followed by the treatment with or without CHX (50 μM) and 25-HC (2.5 μM) for 30 or 90 min as indicated. Insig-1 stabilization was assessed by immunoblot as above. The asterisk denotes a non-specific band.

To further validate the effect of CH25H expression on Insig-1 degradation, we employed the CHO-CH25H^tet-on^ cells to control the levels of CH25H expression. The results showed that the increasing CH25H expression augments the rate of Insig-1 stabilization (**Figure 5C**). These results indicate that endogenous 25-HC inactivates SREBP-2 by protecting Insig-1 from proteasomal degradation.

### Quantitative determination of oxysterols produced by cholesterol hydroxylases

Cholesterol hydroxylases catalyze the addition of a hydroxyl group to cholesterol. However, their specificity is not fully validated quantitatively in living cells. Therefore, we finally sought to determine the ability and specificity of CH25H, CYP27A1, CYP46A1, and CYP7A1 to produce oxysterols. Cellular lipids were extracted from CHO-K1 cells expressing either one of four hydroxylases, and non-saponifiable lipids (containing sterols) were subjected to GC-MS/MS analysis (**Figure 6A, Table S1**). The results show that the expression of CH25H lead to robust production of 25-HC but not of other oxysterols, indicating the high specificity of this enzyme to catalyze cholesterol hydroxylation at the C25 position. CYP27A1 expression predominantly generated 27-HC and small amounts of 25-HC. Approximately only 12.5% of oxysterol synthesized by CYP27A1 was 25-HC. 24S-HC was the major oxysterol produced by CYP46A1 expression. Also, its expression produced small amounts of 25-HC and 27-HC, accounting for 17.9% and 4.8% of oxysterols produced, respectively. CYP7A1 expression specifically produced 7a-HC. On the other hand, the addition of each oxysterol into the culture medium robustly increased cellular amounts of corresponding oxysterols much more than endogenous synthesis by forced expression of each hydroxylase (**Table S1**). 7α,25-dihydroxycholesterol (diHC) and 7α,27-diHC were as low as undetectable levels in CHO-K1 cells.

**Figure 6.**
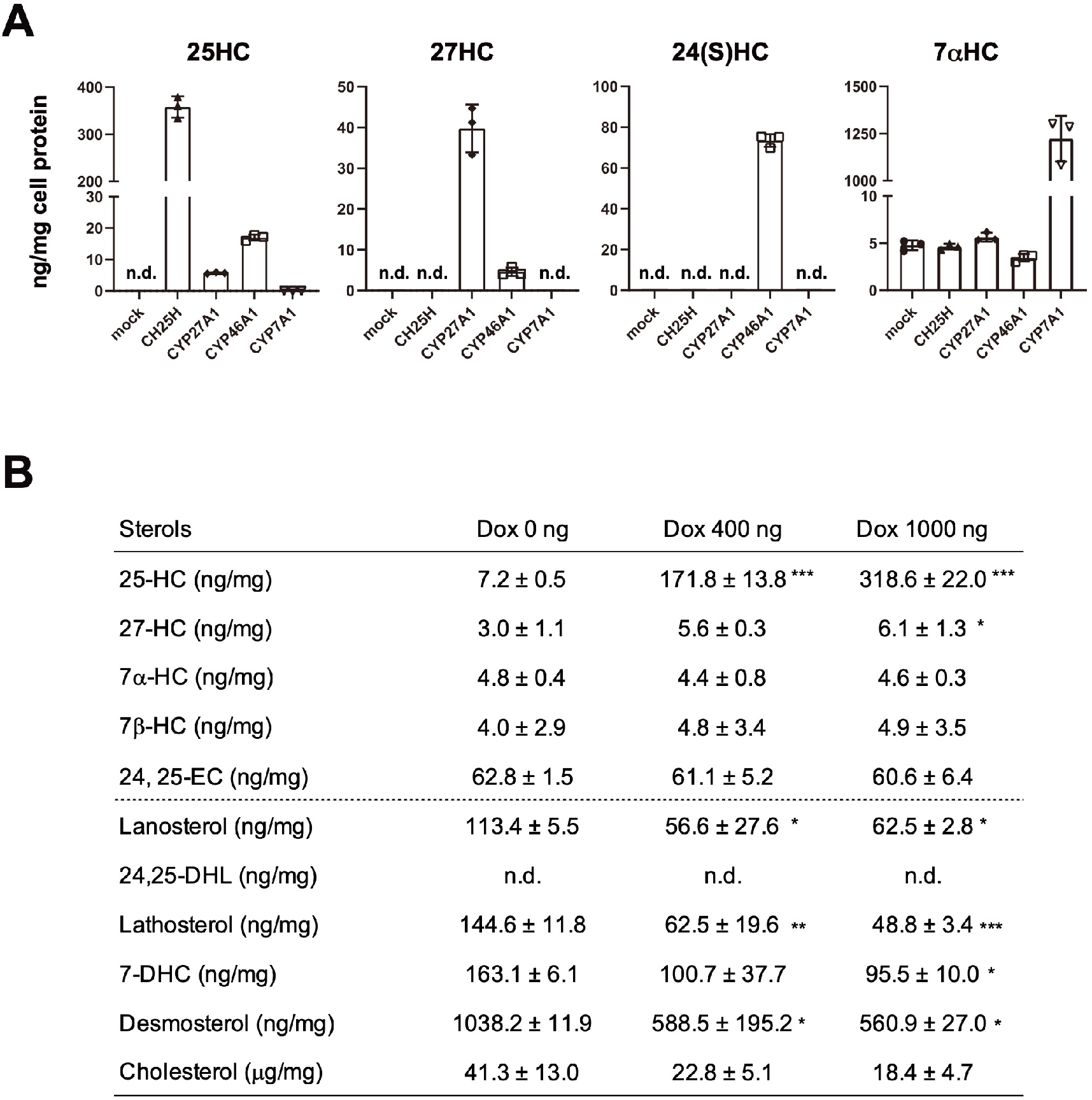
Quantitative determination of oxysterols synthesized by the expression of cholesterol hydroxylases. (A) Production of oxysterols by the expression of CH25H, CYP27A1, CYP46A1, and CYP7A1. FLAG-CH25H, CYP27A1-FLAG, CYP46A1-FLAG, or CYP7A1-FLAG was expressed in CHO-K1 cells. Twenty-four hours after transfection, cellular lipids were extracted and subjected to GC-MS/MS analysis as described in Materials and Methods. The amounts of each sterol were normalized by cell protein. See also Table S1. Data represent means ± S.D. from three biological replicates. (B) Effect of CH25H expression levels on cellular sterol composition. CHO-CH25H^tet-on^ cells were untreated or treated with different concentration of Dox (400 ng/mL or 1000 ng/mL) for 24 hr. Sterol contents were determined by GC-MS/MS. Data represent means ± S.D. from three biological replicates. Statistical analyses were performed by one-way ANOVA with Dunnett post hoc test (*p<0.05, **p<0.01, ***p<0.001). 24,25-DHL, 24, 25-dihydrolanosterol; 7-DHC, 7-dehydrocholesterol.

Next, we validated the CH25H level-dependent production of 25-HC using the CHO-CH25H^tet-on^ cells. The results show that the cellular amounts of 25-HC correlate to the amounts of Dox added (**Figure 6B**). The addition of 0.4 and 1.0 μg/ml of Dox increased cellular 25-HC contents by 24 and 44 times, respectively, compared to cells not treated with Dox. Consistent with Figure 6A, cellular contents of other oxysterols were not altered by CH25H expression, except for a slight increase in 27-HC. The amounts of intermediate sterols were also measured to ensure suppression of SREBP-2 by CH25H expression. The results showed that CH25H expression caused the reduction of intermediate sterols, including lanosterol, lathosterol, 7-dehydrocholesterol, and desmosterol. Thus, it was demonstrated that CH25H expression leads to the production of 25-HC, which then inhibits SREBP-2 activity and cholesterol biosynthesis.

## Discussion

Supraphysiological levels of oxysterols added exogenously can regulate both SREBP-2 and LXR activities in cultured cells (13, 14). However, oxysterols are present in the circulation and tissues at low levels (15, 19). Studies show that oxysterol synthesis is tightly regulated at transcriptional levels (26, 27). Whether oxysterols, particularly side-chain oxysterols, endogenously synthesized within cells provoke the same cholesterol homeostatic responses as exogenous oxysterols remains largely unknown. In this work, we first carefully determined the sensitivity of SREBP-2 and LXR to a series of oxysterols at various concentrations and found that SREBP-2 was much more sensitive to 25-HC and 27-HC than LXR; in the case of 25-HC, 20 – 100 nM was sufficient to inactivate SREBP-2, while for LXR activation, 500 nM or higher concentrations were required. These results indicate that sidechain oxysterols primarily suppress cholesterol synthesis by inactivating SREBP-2 rather than promoting cholesterol efflux by upregulating ABCA1 and ABCG1 expression. On the other hand, at higher concentrations they could activate LXR without adding RXR ligand, confirming that side-chain oxysterols serve as LXR ligands (14). We next examined whether endogenous oxysterols regulate SREBP-2 and LXR by expressing CH25H, CYP27A1, CYP46A1, or CYP7A1. The results revealed that endogenous side-chain oxysterols selectively regulate the SREBP-2 pathway and do not serve as preferential ligands for LXR. This is the first to comprehensively examine the effects of endogenous and exogenous oxysterols on cholesterol homeostatic responses in a single cell system. Our results have uncovered the hydroxylation site-specific and production-dependent roles of oxysterols in cellular cholesterol homeostasis.

How do *de novo* synthesized oxysterols regulate cholesterol homeostatic responses? CH25H and CYP46A1 localize to the ER and synthesize 25-HC and 24S-HC, respectively, in the ER membrane (17), where SREBP, SCAP, and Insig form a trimeric complex (1). CYP27A1 resides in the inner mitochondrial membrane and catalyzes the conversion of cholesterol into 27-HC with the function of StAR (17). The mitochondria often form membrane contact sites with the ER, which facilitates the exchange of lipids between these two organelles (28). Thus, 27-HC produced in the mitochondria may efficiently be transported to the ER. Accordingly, these three side-chain oxysterols could be enriched in the ER membrane and rapidly bind Insig to inhibit its degradation and suppression of the translocation of the SREBP2-SCAP complex to the Golgi (**Figure 7**). On the other hand, the current results have demonstrated that the expression of enzymes catalyzing side-chain hydroxylation of cholesterol and subsequent biosynthesis of corresponding oxysterols do not lead to the activation of LXR, indicating that side-chain oxysterols synthesized in the ER and mitochondria are poorly available for LXR. The majority of LXR localizes in the nucleus. Thus, we hypothesize that endogenous oxysterols hardly enter the nuclei, or other more potent endogenous LXR ligand exists as suggested previously (29). In contrast, exogenous side-chain oxysterols are capable of inducing LXR target gene expression at higher concentrations. Previous studies showed that in addition to the nucleus, LXR localizes to the plasma membrane where it interacts with ABCA1 (30), a prominent LXR target. Oxysterols added exogenously enter cells from the PM. Therefore, they could promptly interact with and activate PM-localized LXR than those synthesized intracellularly. Intracellular distribution and transport of oxysterols remain poorly understood. Further studies are required to clarify whether oxysterols synthesized *de novo* are trapped by a certain oxysterol-binding protein not to enter the nucleus, whether a specific transporting machinery is required for nuclear transport of oxysterols, and whether a more preferential endogenous LXR ligand exists.

**Figure 7.**
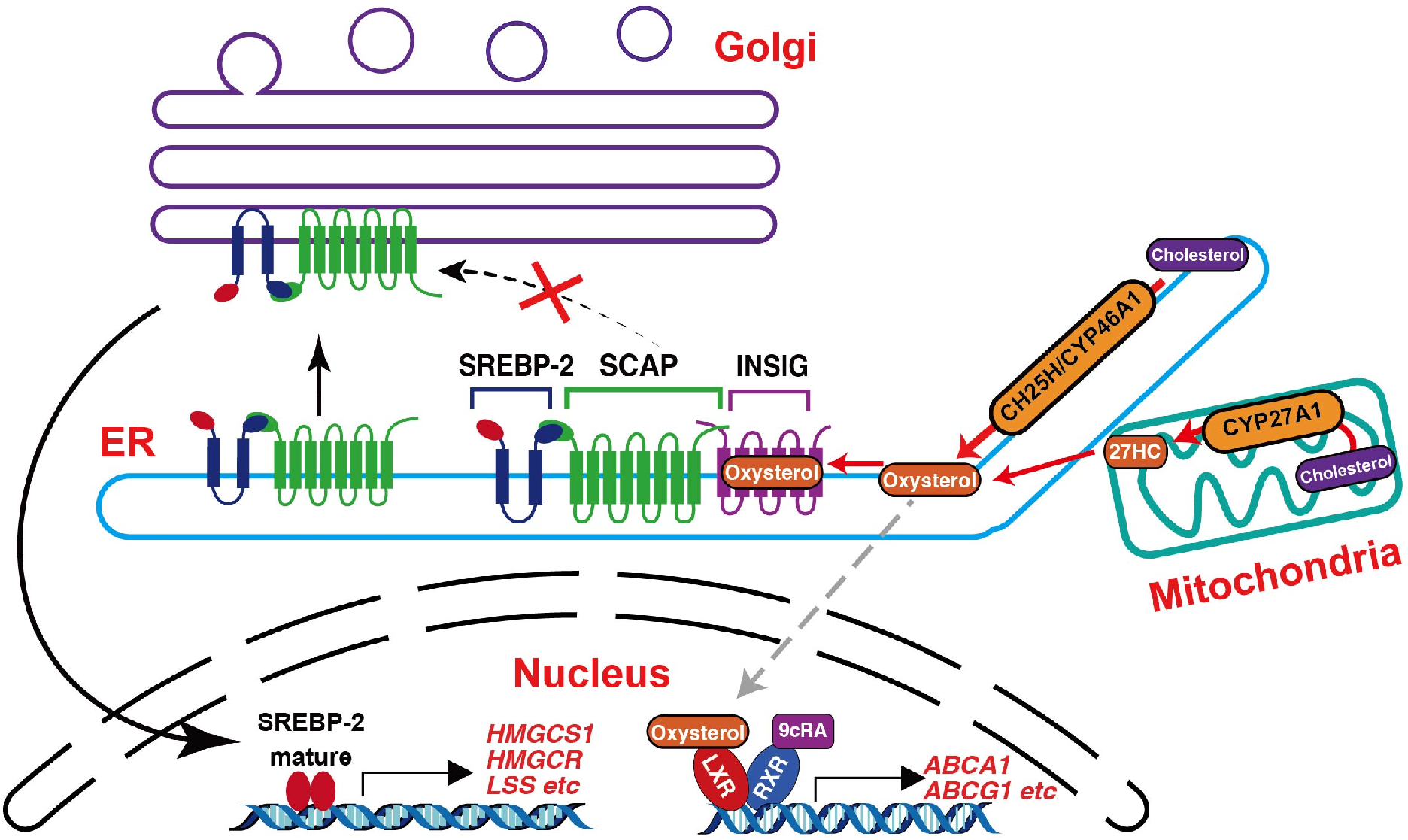
Model of cholesterol homeostatic regulation by endogenous oxysterols. Side-chain oxysterols (25HC, 24S-HC, and 27-HC) synthesized in the ER and mitochondria rapidly bind Insig proteins and inactivate SREBP-2 pathway. Endogenous side-chain oxysterols are poorly available for LXR as its ligand. See text for more details.

In analogy to our findings, whether side-chain oxysterols serve as endogenous LXR ligands remains controversial. In *Ch25h/Cyp27a1/Cyp46a1* triple knockout (TKO) mice, the expression of some LXR target genes in the liver was poorly induced when mice were fed a cholesterol-rich diet (31). However, the effect of the deletion of these three hydroxylase genes on LXR was relatively modest. Furthermore, cholesterol loading of macrophages from TKO mice can markedly up-regulate the expression of LXR target genes (32). Among sidechain oxysterols, 24S-HC is one of the most potent LXR ligands in cultured cells (14, 33). However, overexpression of CYP46A1 in mice showed little effect on LXR target gene expression despite the accumulation of 24S-HC in tissues and blood (34). Similarly, in *CYP27A1* transgenic mice and *Cyp7b1^-/-^* mice where blood and liver 27-HC, and 25-HC and 27-HC markedly increase, respectively, the upregulation of LXR target genes in the liver was undetectable (35). All these results show that the increase in side-chain oxysterols is insufficient to activate LXR *in vivo.* Similar observations were reported for SREBP-2 regulation *in vivo.* Both in *CYP46A1* and *CYP27A1* transgenic mice, SREBP-2 target gene expression is not down-regulated in the liver (34, 35). Furthermore, in *Cyp7b1^-/-^* mouse livers, the expression of SREBP-2 target genes and the rate of cholesterol biosynthesis are largely unaffected (35, 36). These results may support the view that 25-HC, 27-HC, and 24S-HC play only an auxiliary role in cholesterol homeostatic responses in the liver. However, SREBP-2-dependent sterol synthesis is required for the expression of the LXR target gene *Srebp-1c,* but not *Abca1,* in mouse livers (29), suggesting that a certain sterol(s) serves as an endogenous LXR ligand. On the other hand, recent studies have identified CH25H as an infection-inducible gene (26). Lipopolysaccharide and interferon markedly increase CH25H expression and 25-HC production in immune cells, including macrophages (37–39). CH25H-dependent synthesis of 25-HC protects cells from virus infection by inhibiting replication and fusion and exerts anti-inflammatory responses (26). These effects depend on the suppression of SREBP-2 activity and cholesterol biosynthesis (40, 41), indicating that 25-HC-dependent inactivation of SREBP-2 plays a crucial role in immune response both *in vitro* and *in vivo.*

In addition to side-chain oxygenated cholesterol (such as 25-HC, 27-HC, and 24S-HC), desmosterol and 24,25-epoxycholesterol (24,25-EC) can serve as LXR ligands (42, 43). Desmosterol is an intermediate sterol, while 24,25-EC is synthesized from di-oxidosqualene by the shunt pathway or directly from desmosterol by CYP46A1 (7, 16). Our findings showed that in CHO cells, CH25H expression reduced cellular desmosterol contents, which is consistent with the suppression of SREBP-2. Furthermore, our data showed that CYP46A1 expression did not increase cellular amounts of 24,25-EC, suggesting that CYP46A1 plays a minor role in the conversion of desmosterol to 24,25-EC in living cells, at least in CHO cells although this enzyme has the 24, 25-epoxidation activity on desmosterol *in vitro* (44). Therefore, it is conceivable that the expression of cholesterol side-chain hydroxylases suppresses SREBP-2 activity and reduces cellular contents of desmosterol and/or 24,25-EC. In line with this view, the HMGCR inhibitors statins suppress LXR activity (45). Our results thus suggest that desmosterol and 24,25-EC may not serve as preferred ligands for LXR in a context where cholesterol side-chain hydroxylase expression is up-regulated. Therefore, we speculate that desmosterol serves as an endogenous LXR ligand when cholesterol biosynthesis is activated. This hypothesis may be supported by our previous findings that ABCA1 releases not only cholesterol but also intermediate sterols such as lanosterol (46).

Since lanosterol is more cytotoxic than cholesterol and is a poor substrate for ACAT1, desmosterol-dependent upregulation of ABCA1 may provide cells with means to eliminate excess cytotoxic intermediate sterols. In addition, it has been shown that desmosterol accumulates in macrophage foam cells within atherosclerotic lesions and activates LXR (32).

In conclusion, we disclosed the role of endogenously synthesized side-chain oxysterols in cellular cholesterol homeostasis; they selectively suppress SREBP-2 activity by protecting Insig proteins from their proteasomal degradation without serving as LXR ligands. Moreover, this study is the first to comprehensively and quantitatively identify oxysterols produced by the four major cholesterol hydroxylases, CH25H, CYP27A1, CYP46A1, and CYP7A1, in living cells.

## Materials and Methods

### Materials

Sterols were obtained from commercial sources as follows; cholesterol (C8667), 24(S)-hydroxycholesterol (SML1648), 25-hydroxycholesterol (H1015), 27-hydroxycholesterol (SML2042), 7α,25-Dihydroxycholesterol (SML0541), 7-dehydrocholesterol (30800), and 7β-hydroxycholesterol (H6891) from Sigma; desmosterol (NS460402), lanosterol (NS460102), lathosterol (NS460502), and 24,25-dihydrolanosterol (NS460201) from Nagara Science; 7a,27-dihydroxycholesterol (700024P), 7a-hydroxycholesterol (700034P), cholesterol-d7 (700172), 25-hydroxycholesterol-d6 (LM4113-1EA) from AVANTI; 24,25-epoxycholesterol (Ab141633) from Abcam. T0901317 was obtained from Sigma.

### Cell culture

CHO-7 and SRD-15 cells (23) were isolated in the laboratories of Drs. Joseph Goldstein and Michael Brown (UT Southwestern Medical Center), and Dr. Russell DeBose-Boyd (UT Southwestern Medical Center, USA), respectively. CHO-K1 and 25RA cells (47) were kind gifts of Dr. Ta-Yuan Chang (Geisel School of Medicine at Dartmouth, USA). All the CHO cell lines were maintained in DMEM/F12 1:1 mixture supplemented with 7.5% FBS and penicillin/streptomycin. A2058 (obtained from JCRB Cell Bank) and HEK293T cells were cultured in DMEM with 10% FBS and penicillin/streptomycin. All cells were grown in a humidified incubator at 37°C with 5% CO2.

### Plasmid constructs

The coding sequence of human *CH25H* was amplified from human genomic DNA (because human *CH25H* does not contain any introns) and cloned into p3×FLAG-CMV-10 expression vector (Sigma) at the BamHI/EcoRI site to generate pFLAG-CH25H. pFLAG-CH25H was used as a template to create the active site-mutant CH25H (CH25H-H242Q/H243Q) expression plasmid. The 3×FLAG-CH25H sequence was amplified using pFLAG-CH25H as a template and cloned into pTetOne vector (Clontech) at the BamHI/MluI site to generate pFLAG-CH25H^Tet-On^. The coding sequence of human *CYP27A1, CYP7A1,* and *CYP46A1* was amplified by PCR and cloned into p3×FLAG-CMV-14 expression vector (Sigma) at EcoRV/Xbal, HindIII/BamHI, and EcoRI/ BamHI site, respectively. These CYP27A1-FLAG, CYP7A1-FLAG, and CYP46A1-FLAG expression constructs were referred to as pCYP27A1-FLAG, pCYP7A1-FLAG, and pCYP46A1-FLAG, respectively. cDNA for human StAR was cloned into p3×FLAG-CMV-14 expression vector as described above. Primers used for cloning are listed in **Table S2**.

### Transfection and isolation of stable transfectants

Cells were seeded into 6-well plate. Eighteen to twenty-four hours later, cells were transfected with plasmids using Lipofectamine LTX Reagent (Thermo Fisher Scientific) according to a manufacture’s protocol. Twenty-four to forty-eight hours after transfection, cells were used for further experiments.

For the isolation of CHO-K1 cells harboring pFLAG-CH25H^tet-on^ (CHO-CH25H^tet-on^ cells), CHO-K1 cells seeded into 6-well plate were co-transfected with pFLAG-CH25H^tet-on^ (2 μg/well) and pMAM-BSD (0.1 μg/well). Twenty-four hours after transfection, cells were grown in medium containing blasticidin (7 μg/mL) for 5 days. Blasticidin-resistant clones were isolated by limited dilution, and clones that were positive for Dox-dependent FLAG-CH25H expression were selected by immunoblotting with anti-FLAG antibody.

### RNA isolation and quantitative PCR

Total RNA was isolated using ISOGEN II (Nippon Gene). cDNA was then reverse synthesized using total RNA (2 μg/reaction) isolated and High-Capacity cDNA Reverse Transcription Kit (Applied Biosystems). mRNA levels of a gene was analyzed by quantitative real-time PCR (qPCR) using FastStart Universal SYBER Green Maser (ROX) (Roche) and a specific primer set (**Table S3**). qPCR was performed using a Quant Studio 6 Flex Real-Time PCR System or a StepOnePlus instrument (Applied Biosystems). mRNA levels were normalized to *18S* ribosomal RNA levels.

### Luciferase reporter assay

CHO-K1 cells were plated into 12-well plate and grown for overnight. On day 1, pGL2 containing hamster *Hmgcs1* promoter (0.25 μg/well), pCMV-β-galactosidase (0.25 μg/well) (for normalization), and plasmids of interest (0.5 μg/well) were transfected using Lipofectamine LTX (Thermo Fisher Scientific). Five hours after transfection, medium was switched to medium supplemented with 0.1% FBS, cells were further incubated for 16 hr. Afterward, cells were lysed with lysis buffer (25 mM Tris-phosphate (pH7.8), 2 mM DTT, 2 mM EDTA, 10% (v/v) glycerol, and 1% (v/v) Triton-X100), and cell lysate was subjected to luciferase assay. The luciferase activity was measured using a Lumat LB9508 luminometer (Berthold). β-galactosidase activity was also measured using a SpectraMax M2e (Molecular Devices) at 415 nm for normalization.

### Immunoblotting

Cell lysate was prepared using Urea buffer (50 mM Tris-HCl pH 8.0, 50 mM Na-phosphate pH 8.0, 100 mM NaCl, 8 M urea, 0.5% Protease inhibitor cocktail (Nacalai)) or RIPA buffer (50 mM Tris-HCl pH 7.4, 150 mM NaCl, 1 mM EDTA, 1% NP-40, 0.25% sodium deoxycholate, 0.5% Protease inhibitor cocktail) as described (48). After determination of protein concentration using BCA assay (Thermo Fisher), equal amounts of proteins were subjected to SDS-PAGE and immunoblot analysis. Primary antibodies used are as follows; anti-Insig1 polyclonal antibody (PAB8786) from Abnova, anti-FLAG monoclonal antibody (clone M2, F3165) and anti-β-actin monoclonal antibody (clone AC15, A1978) from Sigma, anti-ABCA1 polyclonal antisera (gift from Dr. Shinji Yokoyama, Chubu University, Japan), anti-SREBP-2 monoclonal antibody (clone 7D4) hybridoma supernatant and anti-HMGCR monoclonal antibody (clone A-9) hybridoma supernatant (gift from Dr. Ta-Yuan Chang, Geisel School of Medicine at Dartmouth, NH). Protein expression was normalized to β-actin level unless specified in figure legends. Band intensities were quantified using Image J software or Evolution Capt software (Vilber).

### Immunostaining and Immunofluorescence microscopy

Cells were seeded onto a 18 mm × 18 mm glass coverslip (Matsunami) placed in a 6-well plate and grown overnight. Cells were then transfected with plasmid as specified in figure legends. Afterward, cells were fixed with 4% paraformaldehyde in PBS for 10 min and permeabilized with 0.1% Triton X-100 for 5 min at room temperature (49). After blocking with 5% FBS in PBS for 1 h, cells were incubated with primary antibodies (anti-Tom20 monoclonal antibody from Cell Signaling Technology, 42406, anti-Calnexin monoclonal antibody from Cell Signaling Technology, 2679, or anti-FLAG monoclonal antibody from Sigma, F3165) for 1 hr. After washing with 1% FBS in PBS three times, specimens were incubated with Alexa Fluor 488-conjugated anti-mouse IgG (Invitrogen, A11029) and Alexa Fluor 568-conjugated anti-rabbit IgG (Invitrogen, A11036) (1:800 dilution) in 2% FBS in PBS for 45 min. Nuclei were stained with DAPI. Specimens were washed with PBS for three times and mounted with ProLong Diamond Antifade Mountant (ThermoFisher). Immunofluorescence confocal images were acquired using a Zeiss LSM800 with Airyscan equipped with a Plan-Apochromat 63×/1.40 Oil DIC M27 objective (Carl Zeiss). Images were processed with a Zen software (Carl Zeiss) and Fiji.

### GC-MS/MS analysis

Cellular lipids were extracted with hexane/2-propanol (3:2, v/v) containing 0.01% dibutyl hydroxytoluene and collected into a glass tube. Cholesterol-d7 (100 ng/tube) and 25-hydroxycholesterol-d6 (10 ng/tube) were added to the sample as internal controls. After evaporation under nitrogen gas stream, 1 mL of 100% EtOH and 300 μL of 10 N KOH (in 75% EtOH, v/v) was added to each tube, and lipids were saponified at 80°C for 1 hr. Afterward, non-saponified lipids were extracted with chloroform and dried under nitrogen gas. The sample was then derivatized in 100 μL of 1:1 pyridine/N-Methyl-N-trimethylsilyl trifluoroacetamide (MSTFA) (GL Science) at 80°C for 1 hr and transferred to a new glass vial for GC-MS/MS analysis. Cellular protein was extracted by 0.1 N NaOH as described.

GC-MS/MS analysis was performed using GCMS-TQ8040 NX (Shimadzu) equipped with an AOC-20i autosampler (Shimadzu) and a BPX5 GC column (30 m × 0.25 mm, 0.25 μm, TRAJAN). One μL of sample was injected using the autosampler in splitless mode. The temperature of the sample injection port was set at 275°C, and helium was flowed at 49.5 mL/min as carrier gas. The oven temperature was set at 200°C and kept for 1 min. Afterward, the temperature was increased by 25°C/min to 250°C, 15°C/min to 290°C, and 5°C/min to 320°C, and 320°C was kept for 2 min. Detection by MS/MS was performed using monitoring reaction mode. The retention time, quantification ion, confirmation ion, and collision energy for each sterol were summarized in **Table S4**. Cholesterol-d7 and 25-hydroxycholesterol-d6 were used as internal controls for non-hydroxylated sterols and hydroxylated sterols, respectively. Sterol contents were normalized to cell protein.

### Statistical analysis

Data are presented as mean ± SD. Statistical analyses were performed using one-way ANOVA with Dunnett post hoc test with a RStudio software. *P* < 0.05 was considered statistically significant.

## Acknowledgement

We thank Drs. Shinji Yokoyama (Chubu University, Japan) and Ta-Yuan Chang (Geisel School of Medicine at Dartmouth, NH) for antibodies. We also thank Hikari Ozawa (University of Tokyo) for plasmids, and Drs. Yuichi Watanabe and Yu Takahashi (University of Tokyo) for comments and discussion. This work was supported by AMED-CREST grants 20gm091008h and 21gm091008h (to Y.Y. and R.S.) from Japan Agency for Medical Research and Development, KAKENHI grants 19H02908 and 22H02281 (to Y.Y.) and 20H00408 (to R.S.) from the Japan Society for the Promotion of Science, and a research grant from Asahi Group Foundation (to Y.Y.). H.S. was supported by the Japan Society for the Promotion of Science Research Fellowship for Young Scientists.

