## Supplemental Information for "Hydroxylation site-specific and production-dependent roles of endogenous oxysterols in cellular cholesterol homeostasis"

**Supplemental tables: Table S1 – S4**
**Supplemental figures: Figure S1 – S4**

**Table S1**

Oxysterol contents in CHO-K1 cells ectopically expressed the indicated hydroxylases or incubated with the indicated oxysterols for 16 hr.

| Sterols | mock | CH25H | CYP27A1 | CYP46A1 | CYP7A1 | 25HC | 27HC | 24(S)HC | 7αHC |
|---|---|---|---|---|---|---|---|---|---|
| 25HC (ng/mg) | n.d. | 357.8 ± 18.3 | 5.7 ± 0.2 | 17.0 ± 0.8 | 0.7 ± 0.9 | 780.0 ± 281.0 | 3.3 ± 0.5 | 1.1 ± 0.2 | 3.3 ± 0.4 |
| 27HC (ng/mg) | n.d. | n.d. | 39.8 ± 4.8 | 4.6 ± 0.8 | n.d. | n.d. | 350.6 ± 75.8 | 6.3 ± 0.2 | 5.0 ± 0.2 |
| 24(S)HC (ng/mg) | n.d. | n.d. | n.d. | 73.5 ± 2.6 | n.d. | n.d. | n.d. | 600.7 ± 38.2 | 2.1 ± 0.2 |
| 7αHC (ng/mg) | 4.8 ± 0.4 | 4.6 ± 0.3 | 5.6 ± 0.4 | 3.5 ± 0.3 | 1223.8 ± 99.0 | 4.4 ± 0.8 | 3.9 ± 0.2 | 3.3 ± 0.4 | 841.0 ± 90.1 |
| 24, 25-EC (ng/mg) | 41.0 ± 1.4 | 52.9 ± 8.7 | 40.2 ± 2.9 | 37.5 ± 3.0 | 17.3 ± 0.7 | 25.6 ± 0.3 | 32.2 ± 0.9 | 36.9 ± 2.0 | 57.3 ± 1.6 |

**Table S2**

Primers used for cloning

| | |
|---|---|
| pFLAG-CH25H | FW: 5’-CCGAATTCAAGCTGCCACAACTGCTCCG-3’<br>RV: 5’-CCGGGATCCTCACCGCGCTGGGACAGATGCAGT-3’ |
| pCYP27A1-FLAG | FW: 5’-ATATATGATATCGATGGCTGCGCTGGGCTGCGC-3’<br>RV: 5’-TATATATCTAGAGCACTGTCTCTGCAGGAACTGCA-3’ |
| pCYP7A1-FLAG | FW: 5’-ATATATAAGCTTATGATGACCACATCTTTGATTTGGG-3’<br>RV: 5’-ATATATGGATCCCAAATGCTTGAATTTATATTTAAATTCAAT-3’ |
| pCYP46A1-FLAG | FW: 5’-CGGAATTCATGAGCCCCGGGCTGCTGCTGCT-3’<br>RV: 5’-CGGGATCCGCAGGGGGGTGGTGGGGGTGCGGGCT-3’ |
| pStAR-FLAG | FW: 5’-ATATATAAGCTTATGCTGCTAGCGACATTCAAGCTGTGCGCT-3’<br>RV: 5’-ATATATGAATTCCGACACCTGGCTTCAGAGGCAGGG-3’ |
| FLAG-CH25H$^{tet\text{-}on}$ | FW: 5’-ATATATATACGCGTATGGACTACAAAGAC-3’<br>RV: 5’-ATATATGGATCCTCACTACCGCGCTG-3’ |
| FLAG-CH25H$^{H242Q/H243Q}$ | FW: 5’-CAACAGGACCTGCATCACTCTCA-3’<br>RV: 5’-CACCACACCCCCGTACCACCC-3’ |

**Table S3**

Primers used for qPCR analysis

| | |
|---|---|
| Hamster *18s rRNA* | FW: 5’-TAAGTCCCTGCCCTTTGTACACA-3’<br>RV: 5’-GATCCGAGGGCCTCACTAAAC-3’ |
| Hamster *Abca1* | FW: 5’-GCTCTGGTGTTCAGCCTAAT -3’<br>RV: 5’-CTGGTTAGAGCATTCAGGAGTT-3’ |
| Hamster *Abcg1* | FW: 5’-GGGATCAGAACAGTCGCCTG-3’<br>RV: 5’-CGAGGTCTCTCTTATAGTCAGCGTC-3’ |
| Hamster *Hmgcs1* | FW: 5’-CCTATGACTGCATTGGGCG-3’<br>RV: 5’-CCCAGACTCCTCAAACAGCTG-3’ |
| Hamster *Hmgcr* | FW: 5’-CCCAAAGAAAGCTCCAGACA-3’<br>RV: 5’-CACTCTCAGTTTCCACCACTAAT-3’ |
| Hamster *Sqs* | FW: 5’-CCCAAGTCCAGTTCTCATCTAC-3’<br>RV: 5’-CCTTCAGGTGGTCAGGTATTT-3’ |
| Hamster *Lss* | FW: 5’-GGAGCTCTATGTGGAGGACTAT-3’<br>RV: 5’-CTCAGGCTGGTACTATGGAAAC-3’ |
| Hamster *Insig1* | FW: 5’-ATCAACCACGCCAGTGCCAAAT-3’<br>RV: 5’-CGAATGTCCACCACAAGCCCAAAG-3’ |
| Hamster *Insig2* | FW: 5’-GGGTGGTGCTCTTCTTCATTG-3’<br>RV: 5’-CAGGTGGAAAAAGTGTCACGTTT-3’ |
| Hamster *Fasn* | FW: 5’-CATTCATCAGGCCACCATACT-3’<br>RV: 5’-GTCTTCCCACTGGTACACTTTC-3’ |
| Human *CH25H* | FW: 5’-ATCACCACATACGTGGGCTTT-3’<br>RV: 5’-GTCAGGGTGGATCTTGTAGCG-3’ |
| Human *HMGCS1* | FW: 5’-GACTTGTGCATTCAAACATAGCAA-3’<br>RV: 5’-CTGTAGCAGGGAGTCTTGGTACT -3’ |
| Human *SQS* | FW: 5’-ATGACCATCAGTGTGGAAAAGAAG -3’<br>RV: 5’-CCGCCAGTCTGGTTGGTAA -3’ |
| Human *18S rRNA* | FW: 5’-ACCGCAGCTAGGAATAATGGA -3’<br>RV: 5’-GCCTCAGTTCCGAAAACCA -3’ |

**Table S4**

Detailed parameters for each sterol in GC-MS/MS analysis

| Sterols | Retention time (min) | Quantification | | Confirmation | | | |
|---|---|---|---|---|---|---|---|
| | | MRM transition (m/z) | CE (eV) | MRM transition (m/z) | CE (eV) | MRM transition (m/z) | CE (eV) |
| 7α-Hydroxycholesterol | 8.77 | 456 > 208.3 | 18 | 456 > 119 | 30 | 456 > 95.1 | 33 |
| Cholesterol-d7 | 9.18 | 336 > 121.1 | 18 | 375 > 145.1 | 18 | 336 > 109.2 | 18 |
| Cholesterol | 9.23 | 368 > 145.1 | 21 | 329 > 95.1 | 27 | 329 > 81.1 | 24 |
| Desmosterol | 9.53 | 129 > 73 | 15 | 129 > 57.9 | 30 | 129 > 127.1 | 15 |
| 7-Dehydrocholesterol | 9.56 | 351 > 145.1 | 30 | 351 > 128.1 | 42 | - | |
| Lathosterol | 9.71 | 213 > 157.2 | 12 | 213 > 81.1 | 15 | - | |
| 7β-Hydroxycholesterol | 9.71 | 456 > 233.2 | 18 | 233 > 73.1 | 24 | - | |
| 7α, 25-Dihydroxycholesterol | 10.55 | 131 > 73.1 | 18 | 544 > 73.2 | 39 | - | |
| Lanosterol | 10.70 | 393 > 95.2 | 24 | 393 > 187.4 | 12 | - | |
| 24,25-Epoxycholesterol | 10.78 | 143 > 73.1 | 18 | 129 > 73.1 | 15 | 143 > 128.1 | 18 |
| 24(S)-Hydroxycholesterol | 10.98 | 159 > 69.1 | 9 | 159 > 73.1 | 18 | - | |
| 7α, 27-Dihydroxycholesterol | 10.98 | 103 > 73.1 | 9 | 544 > 233.2 | 33 | - | |
| 25-Hydroxycholesterol-d6 | 11.10 | 137 > 73.2 | 15 | 137 > 58.2 | 30 | - | |
| 25-Hydroxycholesterol | 11.16 | 131 > 73.2 | 9 | 131 > 58.1 | 30 | - | |
| 27-Hydroxycholesterol | 11.64 | 129 > 73.1 | 15 | 456 > 131.2 | 42 | 417 > 69 | 42 |

CE: Collision enegy

**Figure S1**

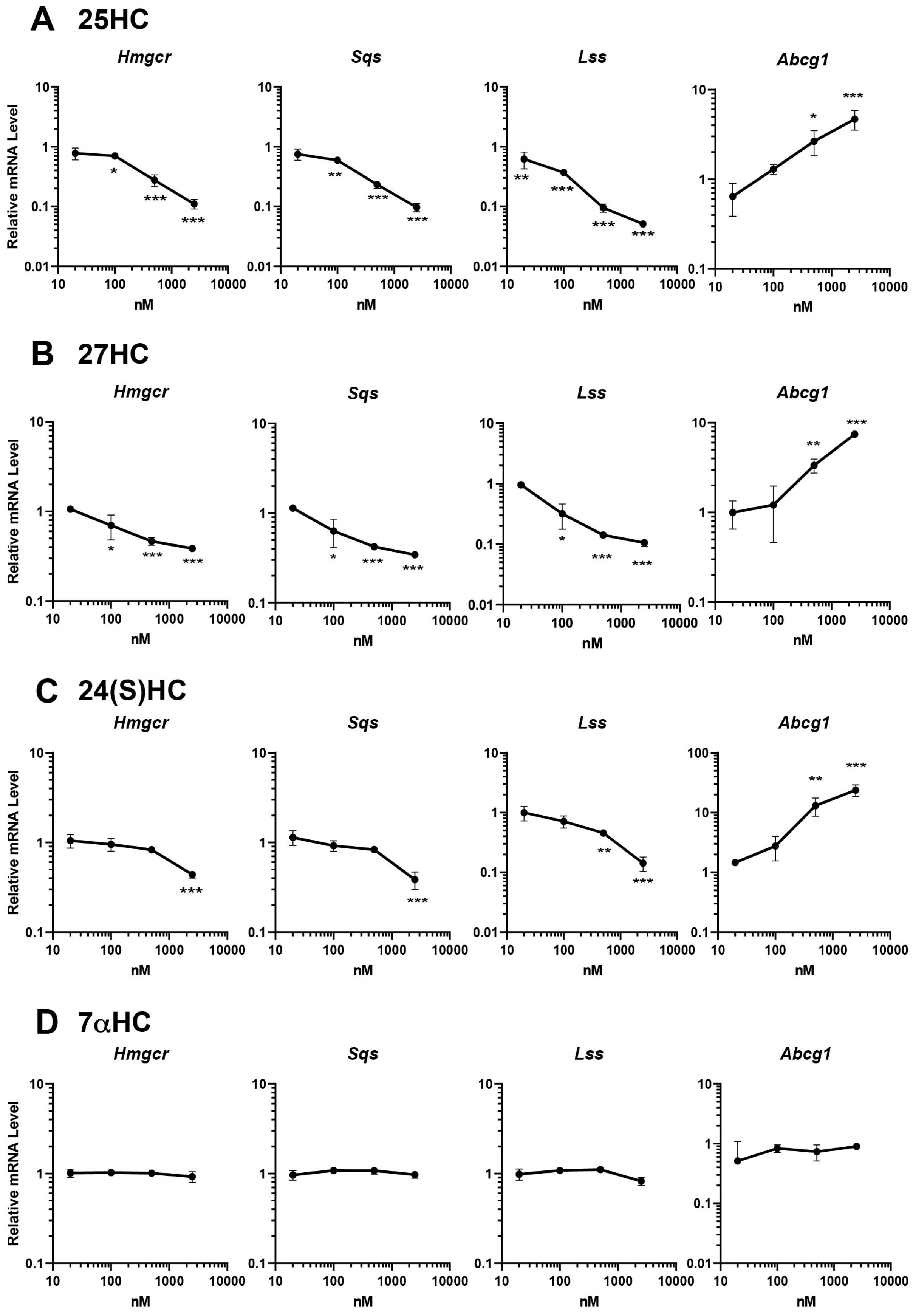

A 25HC
Hmgcr
Sqs
Lss
Abcg1
Relative mRNA Level
nM
*
**
***
B 27HC
Hmgcr
Sqs
Lss
Abcg1
Relative mRNA Level
nM
C 24(S)HC
Hmgcr
Sqs
Lss
Abcg1
Relative mRNA Level
nM
D 7αHC
Hmgcr
Sqs
Lss
Abcg1
Relative mRNA Level
nM
0.01
0.1
1
10
100
10
100
1000
10000

**Figure S1. SREBP-2 is more sensitive to exogenous side-chain oxysterols than LXR, Related to Figure 1.**

(A) CHO-K1 were treated without or with 25-HC (A), 27-HC (B), 24S-HC (C), or 7$\alpha$-HC (D) at different concentration for 24 hr as in Figure 1. mRNA levels of the indicated genes were measured by qPCR. Data represent means ± SD (n = 3). Statistical analyses were performed by one-way ANOVA with Dunnett post hoc test by comparing to the vehicle treatment group (*p<0.05, **p<0.01, ***p<0.001).

Figure S2

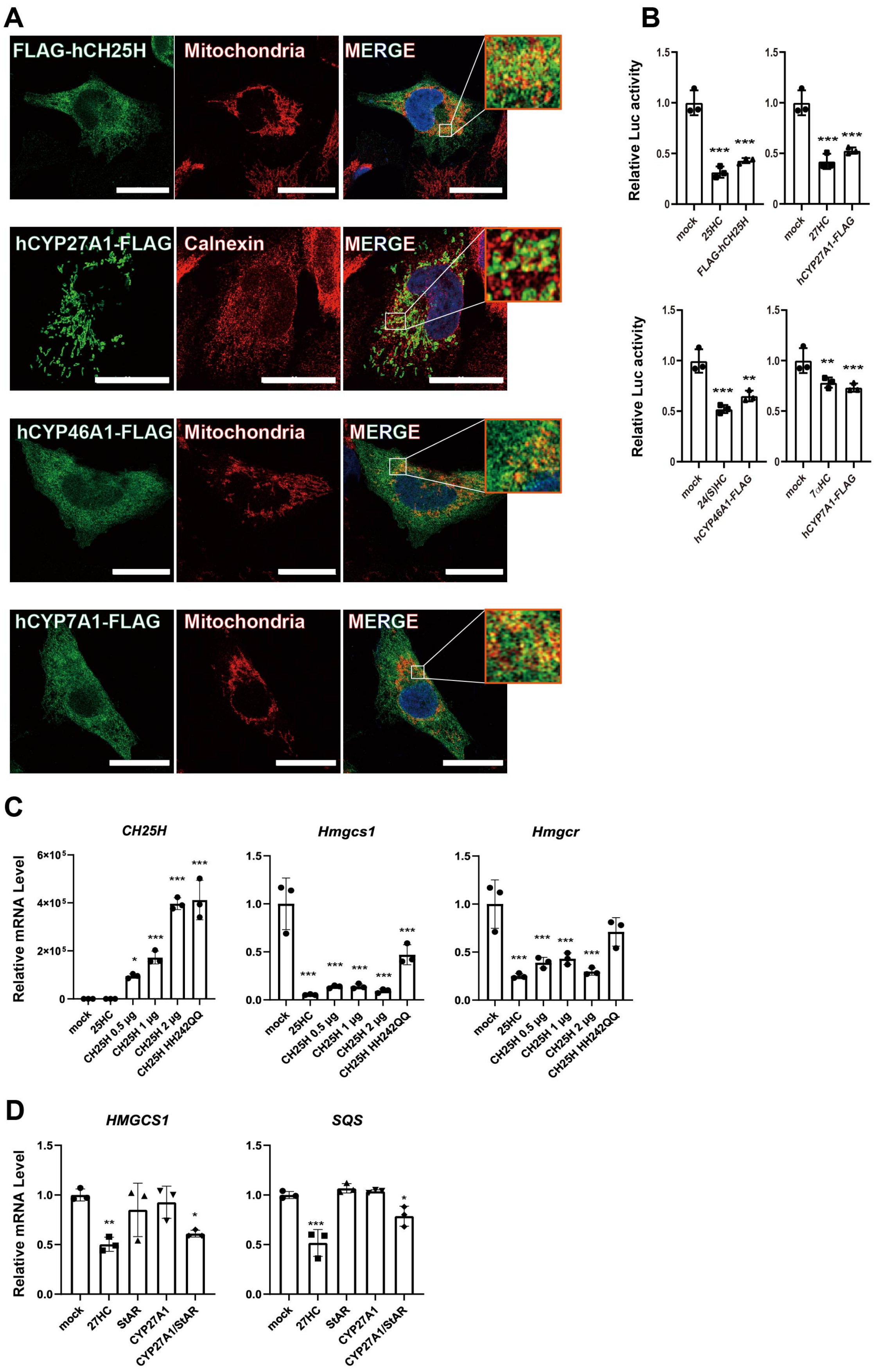
A
FLAG-hCH25H
Mitochondria
MERGE
hCYP27A1-FLAG
Calnexin
MERGE
hCYP46A1-FLAG
Mitochondria
MERGE
hCYP7A1-FLAG
Mitochondria
MERGE
B
Relative Luc activity
mock
25HC
FLAG-hCH25H
mock
27HC
hCYP27A1-FLAG
mock
24(S)HC
hCYP46A1-FLAG
mock
7αHC
hCYP7A1-FLAG
C
CH25H
Hmgcs1
Hmgcr
Relative mRNA Level
mock
25HC
CH25H 0.5 μg
CH25H 1 μg
CH25H 2 μg
CH25H HH242QQ
D
HMGCS1
SQS
Relative mRNA Level
mock
27HC
StAR
CYP27A1
CYP27A1/StAR

**Figure S2. Effect of cholesterol hydroxylase expression of cholesterol homeostatic responses.**

(A) Cellular localization of cholesterol hydroxylases. A2058 cells transfected with the plasmids (pFLAG-CH25H, pCYP27A1-FLAG, pCYP46A1-FLAG, or pCYP7A1-FLAG) were fixed 2 days after transfection. Cholesterol hydroxylases were labeled with anti-FLAG antibody followed by Alexa 488-conjugated anti-mouse IgG (green). The mitochondria and ER were labeled with anti-Tom20 and anti-calnexin antibodies, respectively, followed by Alexa 568-conjugated anti-rabbit IgG (red). Images were taken under a confocal microscopy as described in Materials and Methods. Scale bar, 20 μm except cells with FLAG-CH25H (10 μm).

(B) *Hmgcs1* promoter activity. CHO-K1 cells were transfected with either one of the four hydroxylase expression plasmids (pFLAG-CH25H, pCYP27A1-FLAG, pCYP46A1-FLAG, or pCYP7A1-FLAG) and plasmids for luciferase reporter assay. Mock transfected cells were treated without or with 25-HC, 27-HC, 24S-HC, or 7α-HC (2.5 μM) for 16 hr. Luciferase assay was conducted as described in Materials and Methods.

(C) Effect of mutant CH25H on SREBP-2 target gene expression. CHO-K1 cells seed into 12-well plates were transfected with pFLAG-CH25H (0.5, 1, or 2 μg/well) or pFLAG-CH25H$^{h242Q/H243Q}$ (2 μg/well ). Five hours after transfection, medium was switched to medium containing 0.1% FBS without or with 2.5 μM 25-HC, followed by incubating cells for 16 hr. The expression of the indicated genes were analyzed by qPCR.

(D) Effect of StAR expression of CYP27A1-dependent repression of SREBP-2 taget gene expression. HEK293T cells were transfected with pCYP27A1-FLAG and/or pStAR-FLAG as indicated and incubated in DMEM with 1% FBS in the absence or presence of 27-HC (2.5 μM) for 16 hr. mRNA levels of *HMGCS1* and *SQS* were examined by qPCR.

In B, C and D, error bars represent S.D. from three biological replicates. Statistical analyses were performed by one-way ANOVA with Dunnett post hoc test by comparing to mock (*p<0.05, **p<0.01, ***p<0.001).

**Figure S3**

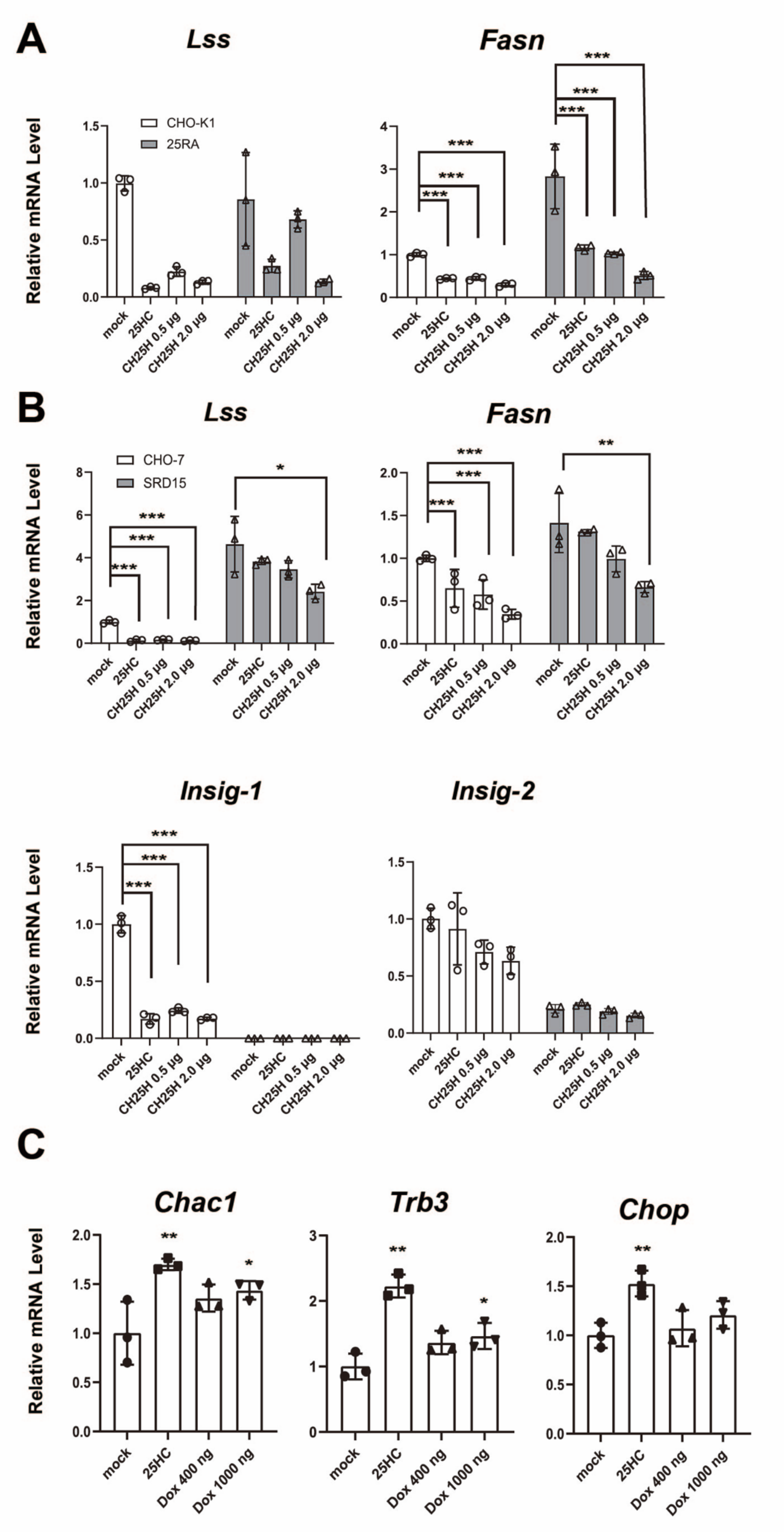

A
Lss
Fasn
CHO-K1
25RA
Relative mRNA Level
mock
25HC
CH25H 0.5 μg
CH25H 2.0 μg
***
B
Lss
Fasn
CHO-7
SRD15
*
**
Insig-1
Insig-2
C
Chac1
Trb3
Chop
Dox 400 ng
Dox 1000 ng

**Figure S3. Endogenous 25-HC regulates cellular responses that required Insig1/2.**

(A, B) Expression of *Lss*, *Fasn*, *Insig-1*, and *Insig-2* in 25RA (A) and SRD-15 (B) cells. Cells were treated as in Figure 4B and C. mRNA levels were analyzed by qPCR. Data represent means ± S.D. from three biological replicates.

(C) Effect of CH25H expression on ATF4 axis. In CHO-hCH25H$^{tet\text{-}on}$ cells, CH25H expression was induced by low (400 ng/mL) or high (1000 ng/mL) concentration of Dox as in Figure 3. The expression of ATF4 target genes (*Chac1*, *Trb3*, and *Chop*) was examined by qPCR.

Statistical analyses were performed by one-way ANOVA with Dunnett post hoc test (*p<0.05, **p<0.01, ***p<0.001).

**Figure S4**

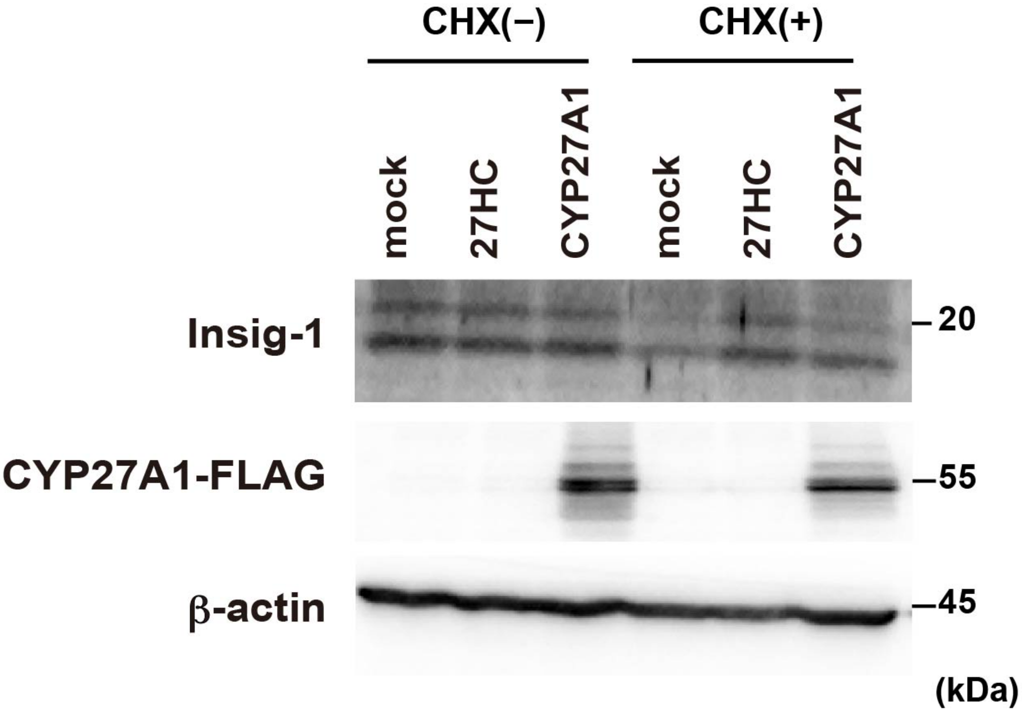


**Figure S4. Stabilization of Insig-1 by CYP27A1 expression.**

CHO-K1 cells transfected with pCYP27A1-FLAG or mock vector were treated without or with CHX (50 μM) ± 27-HC (2.5 μM) for 2 hr as indicated. The expression of Insig-1, CYP27A1-FLAG, β-actin was assessed by immunoblotting. β-actin was used as a loading control.
